# Ancient DNA from Guam and the Peopling of the Pacific

**DOI:** 10.1101/2020.10.14.339135

**Authors:** Irina Pugach, Alexander Hübner, Hsiao-chun Hung, Matthias Meyer, Mike T. Carson, Mark Stoneking

**Affiliations:** Department of Evolutionary Genetics, Max Planck Institute for Evolutionary Anthropology, Leipzig, D04103 Germany; Department of Archaeogenetics, Max Planck Institute for Evolutionary Anthropology, Leipzig, D04103 Germany; Department of Archaeology and Natural History, Australian National University, Canberra, ACT 2061, Australia; Micronesian Area Research Center, University of Guam, Mangilao, Guam, USA

## Abstract

Humans reached the Mariana Islands in the western Pacific by ~3500 years ago, contemporaneous with or even earlier than the initial peopling of Polynesia. They crossed more than 2000 km of open ocean to get there, whereas voyages of similar length did not occur anywhere else until more than 2000 years later. Yet, the settlement of Polynesia has received far more attention than the settlement of the Marianas. There is uncertainty over both the origin of the first colonizers of the Marianas (with different lines of evidence suggesting variously the Philippines, Indonesia, New Guinea, or the Bismarck Archipelago) as well as what, if any, relationship they might have had with the first colonizers of Polynesia. To address these questions, we obtained ancient DNA data from two skeletons from the Ritidian Beach Cave site in northern Guam, dating to ~2200 years ago. Analyses of complete mtDNA genome sequences and genome-wide SNP data strongly support ancestry from the Philippines, in agreement with some interpretations of the linguistic and archaeological evidence, but in contradiction to results based on computer simulations of sea voyaging. We also find a close link between the ancient Guam skeletons and early Lapita individuals from Vanuatu and Tonga, suggesting that the Marianas and Polynesia were colonized from the same source population, and raising the possibility that the Marianas played a role in the eventual settlement of Polynesia.

**Significance Statement:** We know far more about the settlement of Polynesia than we do about the settlement of the Mariana Islands in the western Pacific. There is debate over where people came from to get to the Marianas, with various lines of evidence pointing to the Philippines, Indonesia, New Guinea, or the Bismarck Archipelago, as well as uncertainty over how the ancestors of the present Mariana Islanders, the Chamorro, might be related to Polynesians. We analyzed ancient DNA from Guam, from two skeletons dating to ~2200 years ago, and found that their ancestry is linked to the Philippines. Moreover, they are closely-related to ancient Polynesians from Vanuatu and Tonga, suggesting that the early Mariana Islanders may have been involved in the colonization of Polynesia.

## Introduction

> “Many books have been written about where the Polynesians came from but nobody cares a straw about where the Guamanians came from. And yet it is probable that they can tell at least as much about the peopling of the Pacific as can the Polynesians.”
>
> — -William Howells, *The Pacific Islanders* (1973), p. 248

The human settlement of the Mariana Islands, in western Micronesia, was in some respects more remarkable than the settlement of Polynesia. And yet, as noted in the quote above and by others (1), the settlement of Polynesia has received far more attention than that of the Marianas. Consisting of 15 islands (of which Guam is the largest and southernmost) stretching across some 750 km of sea, the archipelago is located ~2500 km east of the Philippines and ~2200 km north of New Guinea (Figure 1). The earliest archaeological sites date to around 3.5 thousand years ago (kya) (2), and paleoenvironmental evidence suggests even older occupation, starting around 4.3 kya (3). Thus, the first human presence in the Marianas was at least contemporaneous with, and possibly even earlier than, the earliest Lapita sites in Island Melanesia and western Polynesia that date to after 3.3 kya (4) and are associated with the ancestors of Polynesians. However, reaching the Marianas necessitated crossing more than 2000 km of open ocean, whereas voyages of similar length were not accomplished by Polynesian ancestors until they ventured into eastern Polynesia within the past 1000 years (1, 5).

**Figure 1.**
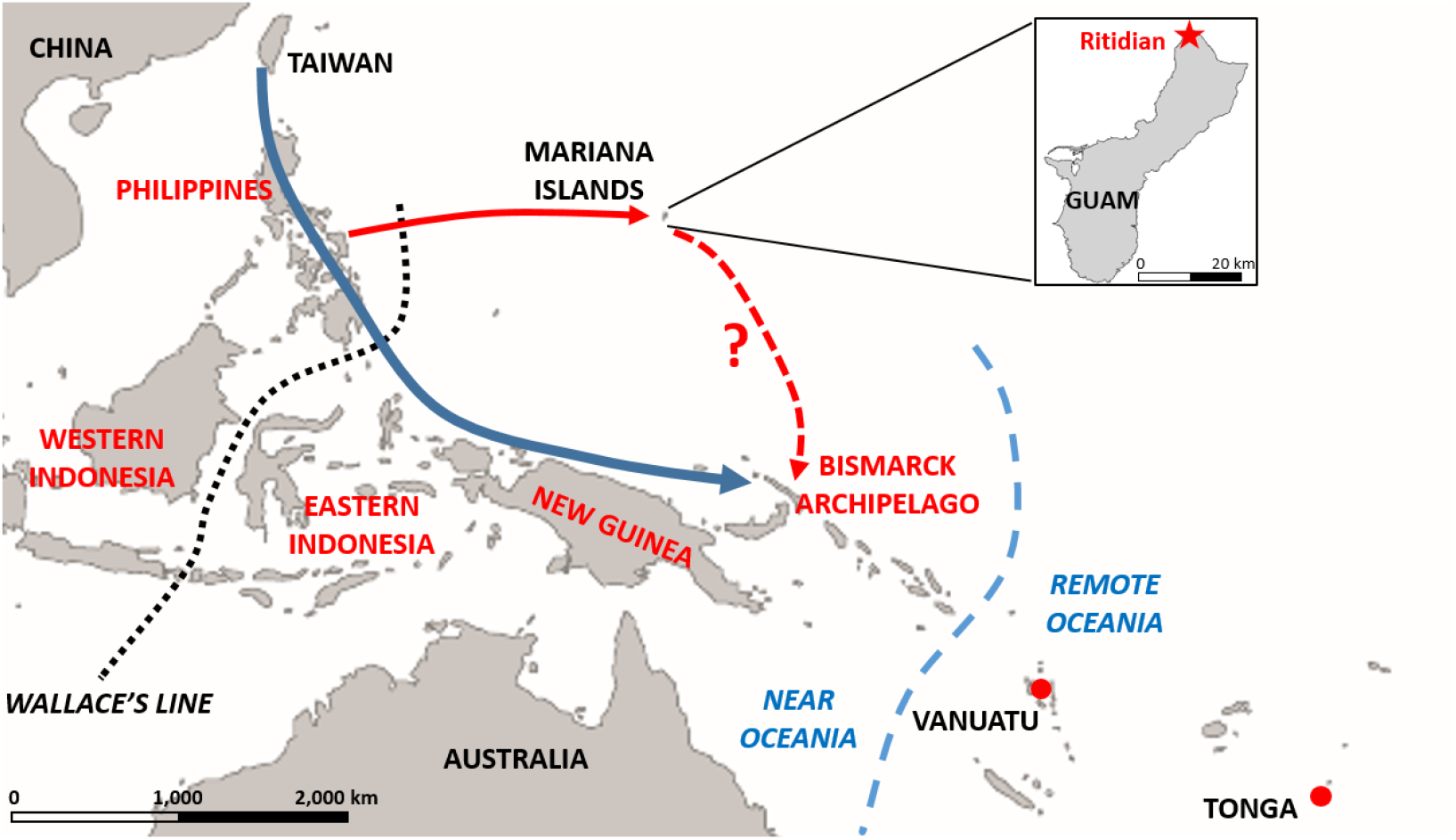
Map of the Western Pacific, showing locations and areas mentioned in the text. Inset shows the location of the Ritidian site on Guam. Location names in red have been suggested as potential sources for the settlement of the Mariana Islands. Wallace’s Line divides biogeographic regions and lies at the boundary of the prehistoric continental landmasses of Sunda and Sahul. The dashed blue line indicates the boundary between Near and Remote Oceania: the islands of Near Oceania were colonized beginning 45-50 kya and involved relatively short, intervisible water crossings, while the islands of Remote Oceania required substantial water crossings that were not intervisible and that were not achieved until ~3.5 kya or later. Red dots indicate the locations of the early Lapita samples from Vanuatu and Tonga; the blue arrow indicates the conventional route for the Austronesian expansion to the Bismarck Archipelago, which was then the source of initial voyages to Remote Oceania; the solid red arrow indicates the route for the settlement of the Marianas supported by this study; and the dashed red arrow indicates the potential contribution of Mariana Islanders to further settlement of the Pacific, suggested by this study.

Where these intrepid voyagers originated from, and how they relate to Polynesians, are open questions. Mariana Islanders are unusual in many respects when compared to othr Micronesians and Polynesians. Chamorro, the indigenous language of Guam, is classified as a Western Malayo-Polynesian language within the Austronesian language family, along with the languages of western Indonesia (the islands west of Wallace’s line; Figure 1), Sulawesi, and the Philippines. Palauan, another indigenous language of western Micronesia, is also a Western Malayo-Polynesian language, whereas all other Micronesian and all Polynesian languages belong to the Oceanic subgroup of Eastern Malayo-Polynesian (6). The most definitive features of Lapita pottery, associated with the earliest presence of Austronesians in Island Melanesia and western Polynesia (7), are absent in the Marianas, as are the domestic animals such as pigs, dogs, and chickens typically associated with Lapita sites and Polynesian settlement (8). Moreover, rice cultivation seems to have been present as an indigenous tradition in the Marianas (9), but so far no such evidence has been found elsewhere in Remote Oceania.

These linguistic and cultural differences have led most scholars to conclude that the settlement of western Micronesia and Polynesia had little to do with one another. To be sure, indications have been noted of morphological (10), cranial (11), and genetic (12–16) affinities between Micronesians and Polynesians, and stylistic links between the pottery of the Philippines, the Marianas, and the Lapita region have also been illustrated (17). Nonetheless, the standard narrative for Polynesian origins (Figure 1) is that they reflect a movement of Austronesian-speaking people from Taiwan beginning 4-5 kya that island-hopped through the Philippines and southeastward through Indonesia, reaching the Bismarck Archipelago around 3.5 – 3.3 kya. From there they spread into Remote Oceania, with subsequent additional migrations from Near Oceania around 2.5 kya that brought more Papuan-related ancestry into Remote Oceania. This narrative is supported by a large body of archaeological, linguistic, and genetic data (7, 18–26), and western Micronesia typically does not figure in this orthodox story.

Compared to Polynesians, the origins of the Mariana islanders are more uncertain. Most mtDNA sequences of modern Chamorros belong to haplogroup E, which occurs across Island Southeast Asia and is thought to be associated with the initial peopling of the Marianas, while the less-frequent haplogroup B4 sequences, which are found in high frequency in Polynesians, are attributed to later contact (27). Studies of a limited number of autosomal short-tandem repeat loci similarly indicate differences in the affinities of western Micronesians (Palau and the Marianas) vs. eastern Micronesians, with the former showing ties to Southeast Asia and the latter to Polynesia (12, 15). The linguistic evidence for Chamorro would suggest an origin from western Indonesia (28) or the Philippines (1), and the oldest decorated pottery and other artifacts of the Marianas, dating to around 3.5 kya, have been matched with counterparts in the Philippines at around the same time or even earlier (29). However, alternative views have been proposed and debated (30, 31), and it is not clear to what extent the genetic and linguistic relationships of the contemporary Chamorro reflect initial settlement vs. later contact. Moreover, computer simulations of sea voyaging found no instances of successful voyaging from the Philippines or western Indonesia across to the Marianas; instead these simulations indicated New Guinea and the Bismarck Archipelago as the most likely starting points (32, 33).

Genomic evidence can shed light on this debate over the origin of the Chamorro, as well as on their relationships with Polynesians. Two main genetic ancestries are present in New Guinea and the Bismarck Archipelago: the aforementioned Austronesian (Malayo-Polynesian), which arrived with the spread of Austronesian speakers from Taiwan, and “Papuan,” which is a general term for the non-Austronesian ancestry that was present in New Guinea and Island Melanesia prior to the arrival of the Austronesians; it should be kept in mind that “Papuan” ancestry is quite heterogeneous in composition across the region (34–36). Papuan-related ancestry probably traces back to the original human populations of the region, at least 49 kya (37), and is readily distinguished from Austronesian ancestry. Papuan-related ancestry is present not only in New Guinea and the Bismarck Archipelago, but also at substantial frequencies in eastern Indonesia (38–40), defined here as all Indonesian islands to the east of Wallace’s line (Figure 1). However, Papuan-related ancestry is practically absent west of Wallace’s Line (Figure 1), so if the first settlers of the Marianas started from the Philippines or west of Wallace’s Line, then they should have had little if any Papuan-related ancestry. Conversely, if they started from eastern Indonesia, New Guinea, or the Bismarck Archipelago, then they should have brought appreciable amounts of Papuan-related ancestry.

In principle, to address this issue, the ancestry of the modern inhabitants of the Marianas could be analyzed for Papuan-related ancestry. However, a common finding of ancient DNA studies is that the ancestry of people in a region today may not reflect the ancestry of people living in that region thousands of years ago (41). In particular for the Marianas, the archaeological evidence indicates substantial cultural change around ~1 kya (27, 42), coinciding with the construction of stone-pillar houses in formal village arrangements *(latte)* at a time when nearly all of the Pacific Islands were populated and connected by long-distance sea voyaging (7). The presence of mtDNA haplogroup B4 sequences in modern Chamorro has been attributed to contact during the *latte* period (27).

In addition, population contacts and movements became more complicated during the European colonial period, starting with the arrival of Magellan in 1521 in the Marianas and continuing with the Manila-Acapulco galleons (and slave trade) from 1565-1815; Guam was a regular stopover on these voyages. European colonialism also involved multiple relocations and reductions in population size across the archipelago. These events undoubtedly had an impact on the genetic ancestry of the modern Chamorros, making it more difficult to assess their origins and potential relationships with Polynesians. It would therefore be preferable to address these issues with ancient DNA from the Marianas.

At the Ritidian Site in northern Guam (Fig. S1), two skeletons clearly pre-dating the *latte* period were found outside a ritual cave site (43). These individuals, RBC1 and RBC2, were buried side-by-side in extended positions, with heads and torsos removed (Fig. S2). Direct radiocarbon dating of a bone from RBC2 produced a result of 2180 +/- 30 years calibrated years bp (43), which is thus some 1000 years after the initial settlement of Guam, but also some 1000 years before the *latte* period. Here we report the analysis of ancient DNA retrieved from these remains; our results contribute to the debate over the starting point for the first voyages that led to human settlement of the Marianas, and we provide additional insights into the role of the Marianas in the larger view of the peopling of the Pacific.

## Results

Shotgun sequencing of libraries constructed from DNA extracted from the ancient Guam skeletons revealed elevated C->T substitutions at the ends of fragments, as expected for ancient DNA (Table S1 and Fig. S3). The percent endogenous DNA was too low for further shotgun sequencing (Table S1); we therefore proceeded by capture enrichment for the mtDNA genome, and for a panel of 1.2 million SNPs used in previous ancient DNA studies (23–25, 44, 45), prior to sequencing.

### MtDNA and Y chromosome

After merging the sequence data from libraries enriched for mtDNA while excluding those that were highly contaminated (Table S2), we were able to obtain mtDNA genome sequences at an average coverage of 95.2-fold for RBC1 and 261.3 fold for RBC2. Estimated contamination in the mtDNA sequences, using a likelihood-based approach (46) subsequently referred to as contamMix (47), was 17.9% for RBC1 and 6.6% for RBC2 (Table S3). The sequences are identical where they overlap, and even with this relatively high level of contamination, both sequences are confidently assigned to haplogroup E2a (Table S3). In addition to the diagnostic mutations for haplogroup E2a, both sequences carry a novel derived substitution at position 8981, which results in an amino acid substitution (Gln → Arg) in the ATP6 gene.

Haplogroup E2a is the most common haplogroup in the modern Chamorro population of Guam (27), with a frequency of 65%. Elsewhere it is reported to occur sporadically in populations from the Philippines and Indonesia (48–51), and in a single individual from the Solomon Islands (52); otherwise, it is absent from Oceania and has not been reported from Mainland Southeast Asia. The finding of this haplogroup in the ancient Guam skeletons thus suggests links to the Philippines and Indonesia, rather than New Guinea or the Bismarck Archipelago. Of additional importance, the high frequency of this haplogroup in modern Chamorros suggests a degree of genetic continuity with the population represented by the ancient skeletons, persisting through the interceding cross-population contacts since the *latte* period after ~ 1 kya and later European colonial events.

Based on the ratio of the average coverage of X chromosome vs. X chromosome + autosomal reads in the shotgun sequencing data, RBC1 is male and RBC2 is female (Fig. S4). The Y chromosome of RBC1 is assigned to haplogroup O2a2 (formerly haplogroup O3a3), based on having the derived allele for the diagnostic marker P201 (53); genotypes at all other informative Y-chromosome SNPs for which there are data from RBC1 are consistent with this haplogroup (Table S4). Haplogroup O2a2-P201 is widespread across Mainland and Island Southeast Asia and Oceania, and has been associated with the Austronesian expansion (54, 55).

### Genome-wide SNP data: ancient Guam origins

We enriched 13 sequencing libraries from RBC1 and 11 from RBC2 for ~1.2 million SNPs (Table S5) and obtained data (Table S6) for 128,772 SNPs (39,760 in deaminated reads) for RBC1 and 361,982 SNPs (143,451 in deaminated reads) for RBC2. Given the relatively high contamination estimates for some of the libraries (Table S5), we either redid analyses using only deaminated reads (if there were enough deaminated reads), or included data from Europeans in the analysis, to ensure that contamination with modern European DNA was not influencing the results. The results reported below are based on all reads, as we did not find any indications of contamination influencing the results.

We first checked if RBC1 and RBC2 might be related by calculating the fraction of pairwise differences for the 33,400 overlapping SNPs between them, and comparing this to mean pairwise distances for first, second, and third degree relatives in the 1000 Genomes Project dataset (56), using sites on the Human Origins Array (Fig. S5A). While the pairwise distance between RBC1 and RBC2 is similar to that for first degree relatives in the 1000 Genomes dataset, suggesting that they might be first degree relatives, we obtained similar mean pairwise distances for other ancient samples from Southeast Asia and Oceania (Fig. S5B). We therefore conclude that the limited amount of data and/or low overall genetic diversity characteristic of the ancient samples preclude accurate assessment of relatedness.

We then projected RBC1 and RBC2 onto a PCA constructed with modern samples genotyped on the Affymetrix 6.0 platform and with data from the Simons Genome Diversity Project (SGDP); the overlap with this array is provided in Table S6 and details on the modern samples are in Table S7. This dataset has good coverage of populations from Island Southeast Asia, in particular from eastern Indonesia, which exhibit both Asian-related and Papuan-related ancestry and hence are a potential proxy for the ancestry in the ancient Guam samples. The results for the first two PCs (Figure 2A) show three axes of variation, with Europe/South Asia, New Guinea, and Southeast Asia at the vertices. The two ancient Guam samples overlap samples from Taiwan and the Philippines. There is no indication of any Papuan-related ancestry in the ancient Guam samples, particularly when compared to eastern Indonesian samples, all of which have some Papuan-related ancestry and hence are clearly separated from other Southeast Asian samples.

**Figure 2.**
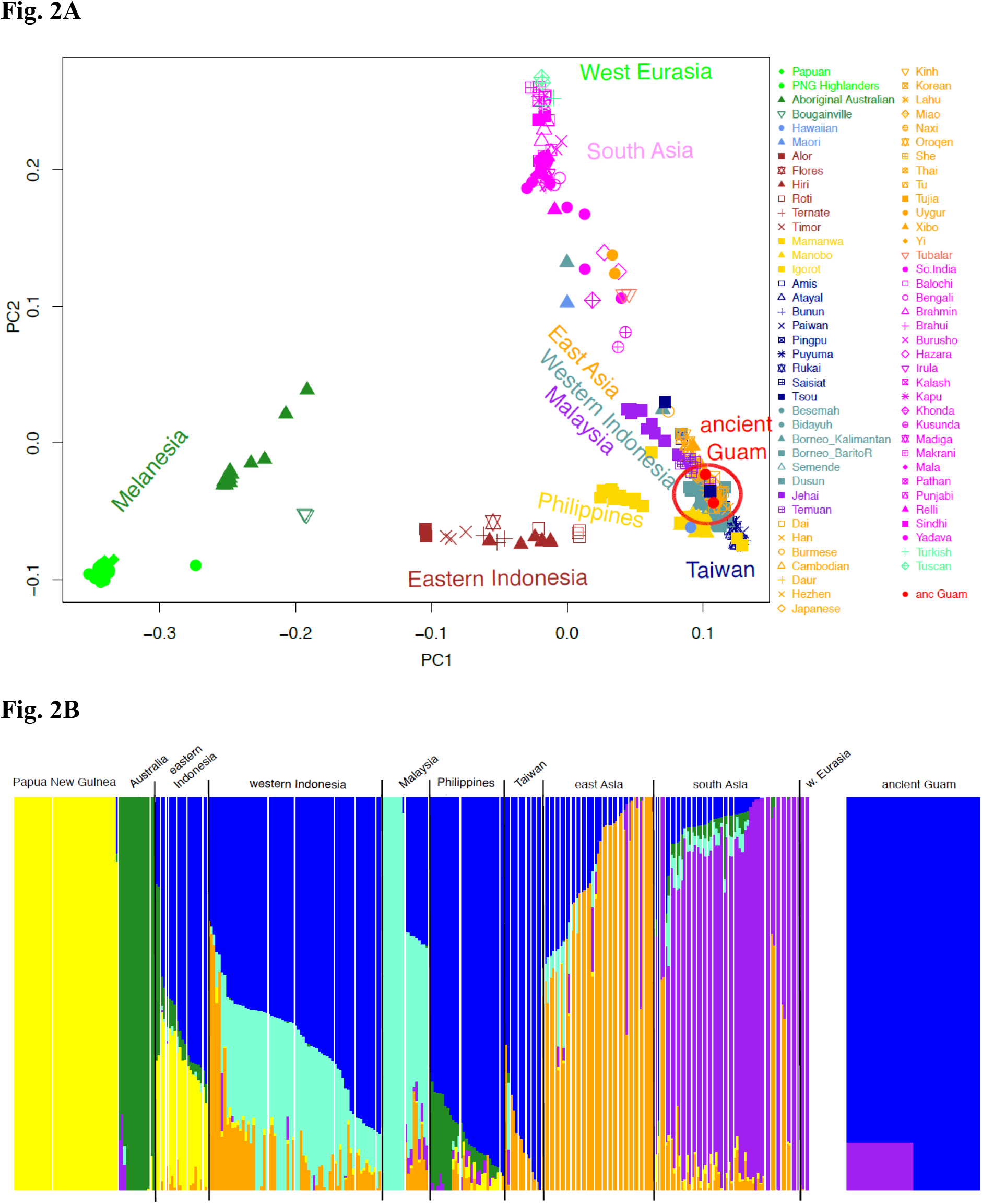
PCA and ADMIXTURE analyses of the ancient Guam samples merged with modern samples genotyped on the Affymetrix 6.0 platform and with SGDP samples. **A**, plot of the first two principal components. The ancient Guam samples are projected. **B**, ADMIXTURE results for K=6.

We next carried out ADMIXTURE analysis of the same dataset; while the results for K=3 are associated with the lowest cross-validation error (Fig. S6A), the results for K=6 distinguish different ancestry components for Mainland vs. Island Southeast Asia, so we show these results in Figure 2B and the results for K=2 to K=8 in Fig. S7. Notably, the yellow ancestry component, which is characteristic of New Guinea and is also present in eastern Indonesia, is completely lacking in the ancient Guam samples for all analyzed values of K (Figure 2B, Fig. S7). Moreover, at K=6 the two ancient Guam samples have the dark blue ancestry component, which is at highest frequency in individuals from the Philippines and Taiwan (Figure 2B). RBC1 also has a purple component which likely reflects recent European DNA contamination.

Thus, these PCA and ADMIXTURE analyses suggest that there is no Papuan-related ancestry in the ancient Guam samples, and moreover indicate that they are most similar to modern samples from the Philippines and Taiwan. However, the number of SNPs in the Affymetrix 6.0 dataset that overlap the ancient Guam samples is too small for more formal tests of population relationships (Table S6), and moreover this dataset has limited coverage of modern Oceanian populations. We therefore carried out all further analyses with the Human Origins dataset, which includes more modern samples from Near and Remote Oceania (Table S7), more overlap with the ancient Guam data (Table S5), and also includes data from ancient samples from Asia and the Pacific (Table S8), including early Lapita samples from Vanuatu and Tonga.

A PCA of these samples with the ancient samples projected (Figure 3A) places the early Lapita samples at one vertex, East Asia at another, and New Guinea at the third vertex; the ancient Guam samples are now projected away from modern Taiwan and Philippine samples, in the direction of the early Lapita samples. An ADMIXTURE analysis of these data for K= 9 (Figure 3B; results for K=5 to K=12 in Fig. S8), which has the lowest crossvalidation error (Fig. S6B), now reveals two primary ancestry components in the ancient Guam samples: a dark blue component as before that is at highest frequency in Indonesia and the Philippines, and an orange component that is at highest frequency in Polynesia; the additional minor purple component likely reflects recent European contamination. As before, there is no indication from either the PCA or the ADMIXTURE analysis of any Papuan-related ancestry in the ancient Guam samples.

**Figure 3.**
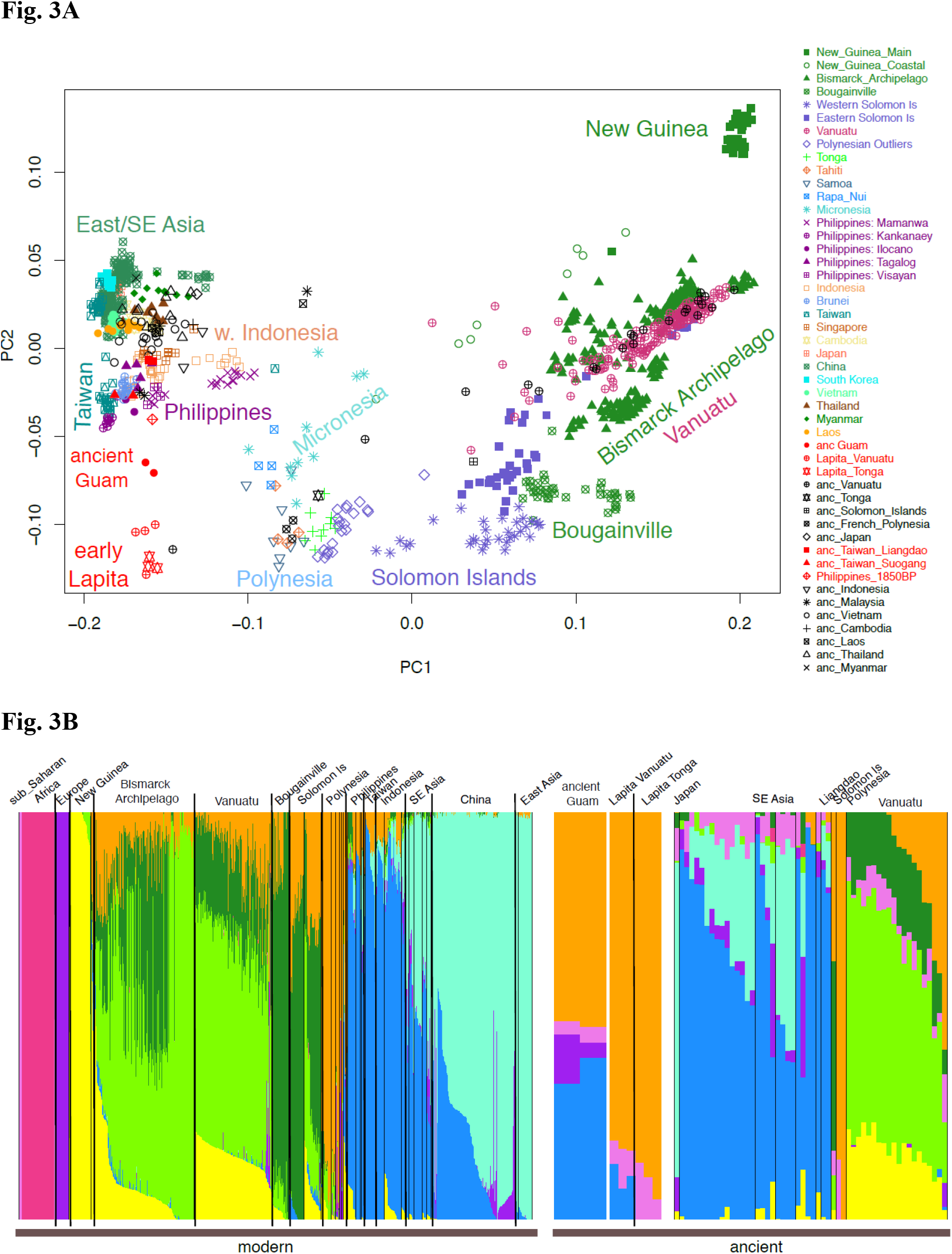
PCA and ADMIXTURE analyses of the ancient Guam samples merged with Human Origins array data for modern and ancient samples. **A**, plot of the first two principal components. Ancient samples are projected. **B**, ADMIXTURE results for K=9.

While the presence of these two ancestry components in the ancient Guam samples could indicate admixture between a source population related to Indonesia/Philippines and another related to Polynesians, other explanations for the presence of multiple ancestry components are possible (36, 57). In particular, it could be that the ancient Guam samples are ancestrally related to both Indonesia/Philippines and to Polynesians, and that subsequent divergence and genetic drift has facilitated the identification of separate Indonesia/Philippine and Polynesia-related ancestry components in the ADMIXTURE analysis, both of which are present in the ancient Guam samples. To investigate the relationships of the ancient Guam samples in more detail, we analyzed outgroup-*f*3 and *f*4 statistics. The outgroup-*f*3 analysis, which compares the amount of drift (i.e., ancestry) shared by the ancient Guam samples with other populations relative to an outgroup (Mbuti), shows that the ancient Guam samples share the most drift with the Lapita Vanuatu and Lapita Tonga samples, followed by an ancient sample from the Philippines and then by modern samples from the Philippines and Taiwan and late Neolithic samples from the Taiwan Strait islands (Figure 4A). Notably, the drift shared with New Guinea, and with French, is less than that with any other population, indicating that the ancient Guam samples show the least relatedness with these two populations. These results further support the lack of any Papuan-related ancestry in the ancient Guam samples, and moreover also indicate that recent European contamination is not influencing these results.

**Figure 4.**
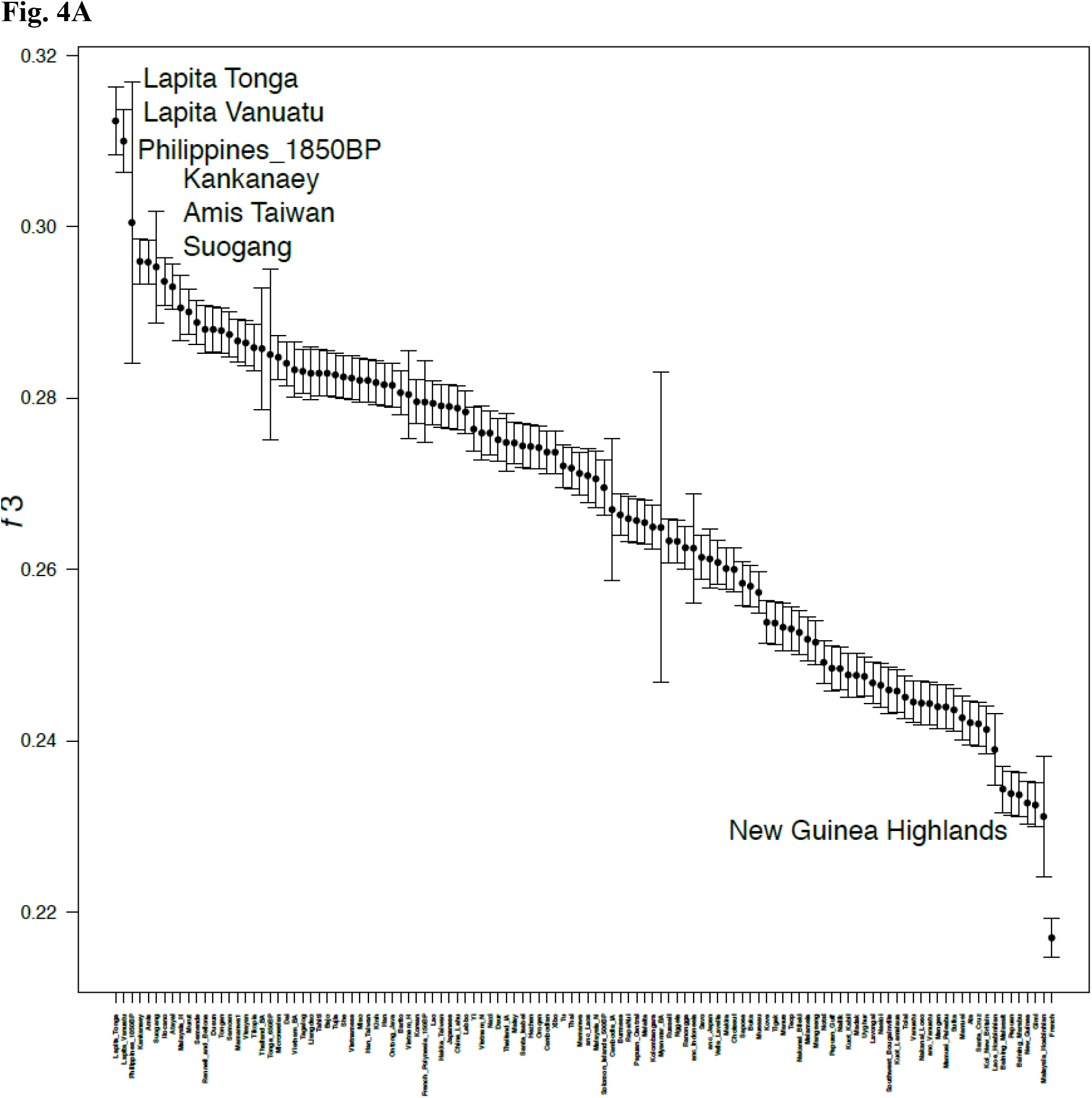
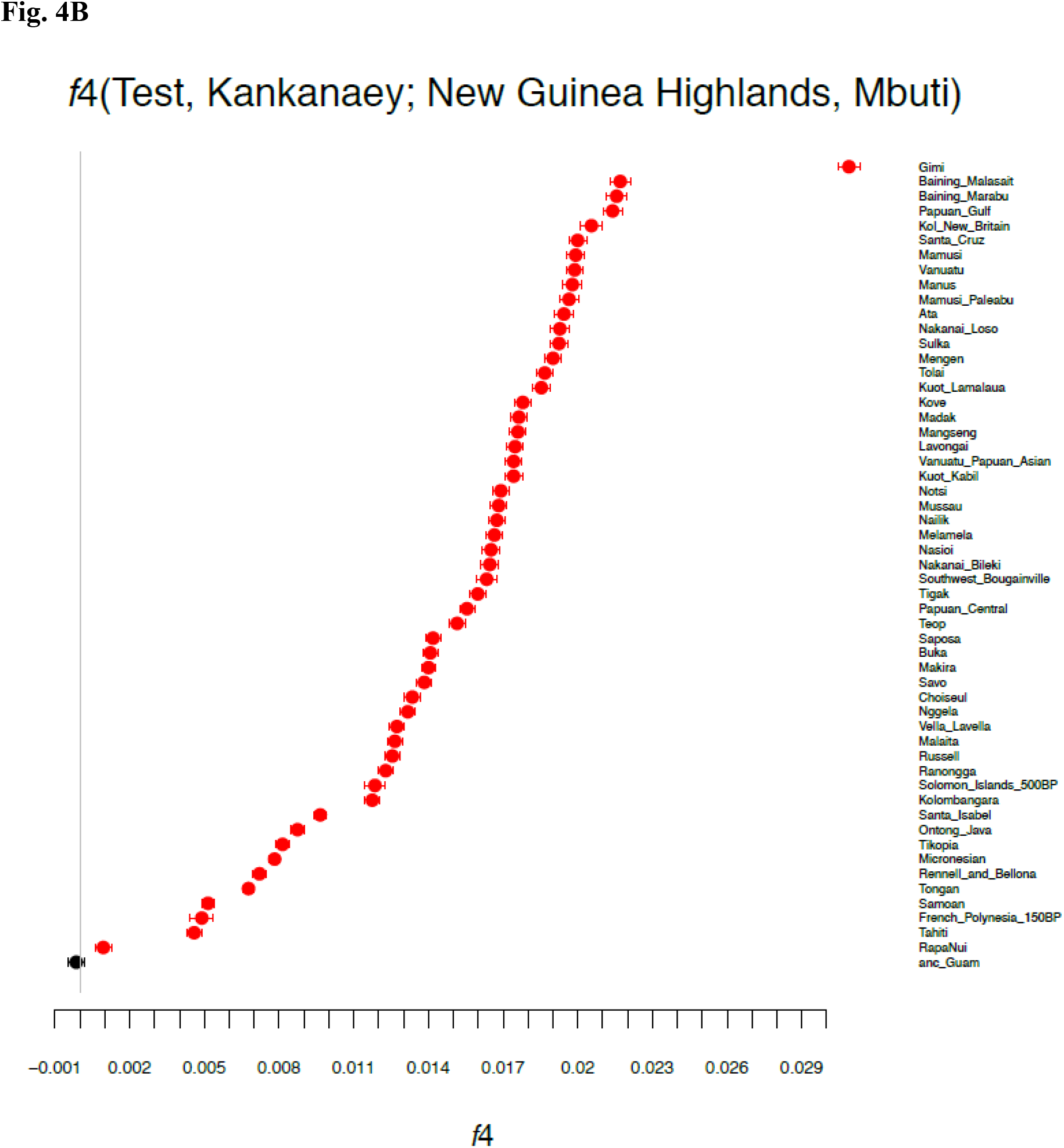
Outgroup-*f*3 and *f*4 results for the relationships of the ancient Guam samples with other populations. **A**, outgroup-*f*3 results comparing the ancient Guam samples to other modern and ancient samples, with Mbuti used as the outgroup. Bars indicate 1 SE. Larger values of the *f3* statistic indicate more shared drift, and hence a closer relationship with the ancient Guam samples. **B**, results for an *f*4 test of the form *f*4(test, Kankanaey; New Guinea highlands, Mbuti). *f*4 values that are significantly different from zero are in red.

We then constructed an *f*4 statistic of the form *f*4(test, Kankanaey; New Guinea highlands, Yoruba); values of this statistic that are equal to zero indicate that the test population forms a clade with Kankanaey relative to New Guinea; values less than zero indicate that Kankanaey shares more ancestry with New Guinea than does the test population; and values greater than zero indicate that the test population shares more ancestry with New Guinea than does Kankanaey. We used the ancient Guam samples and all other populations from Oceania as the test population; the results (Figure 4B) indicate that all populations from Oceania tested share ancestry with New Guinea in comparison to Kankanaey, except for the ancient Guam samples. These form a clade with Kankanaey, as the Z statistic is not significantly different from zero (Z=-1.93)

### Genome-wide SNP data: relationships with early Lapita samples

The PCA, ADMIXTURE, and outgroup-*f*3 analyses not only indicate affinities between the ancient Guam samples and Philippine/Taiwan populations, but additionally suggest strong affinities between the ancient Guam and early Lapita samples. To investigate in more detail the relationships among the ancient Guam and early Lapita samples with samples from Asia and Oceania, we conducted *f*4 analyses of the form (ancient Guam, early Lapita; Asia/Oceania, Mbuti), separately for the early Lapita Vanuatu and Tonga samples and for all modern and ancient Asian and Oceanian samples in the dataset. Values of this *f*4 statistic that are consistent with zero imply that the ancient Guam and early Lapita samples form a clade; negative values indicate excess shared ancestry between the early Lapita sample and the Asia/Oceania population; and positive values indicate excess shared ancestry between the ancient Guam samples and the Asia/Oceania population. The results (Fig. S9) show that the ancient Guam and early Lapita samples always form a clade with one another when compared to any Asian population. However, both of the early Lapita samples share more ancestry with ancient and modern Polynesian samples (but not with any other samples from Oceania) than do the ancient Guam samples. This is further supported by outgroup-*f*3 comparisons of the ancient Guam and early Lapita samples with other populations (Fig. S10): both early Lapita samples share more drift with modern and ancient Remote Oceanians than do the ancient Guam samples. Nonetheless, *f*4 statistics of the form (Oceania, early Lapita; ancient Guam, Mbuti) are always significantly negative for both early Lapita samples, regardless of which Oceanian population is included in the test (Fig. S11). These *f*4 results indicate that there is shared drift between the early Lapita and ancient Guam samples when compared to any other Oceanian sample, in keeping with the outgroup *f*3 results (Figure 4). Overall, the *f*3 and *f*4 results imply that while the early Lapita and ancient Guam samples are closely related to each other, the early Lapita samples are a better source for the Polynesian-related ancestry in modern and ancient Oceanian samples than are the ancient Guam samples.

We next used admixture graphs (i.e., trees that allow for admixture or migration) to further investigate the relationships among the ancient Guam, early Lapita, and other Asian and Oceanian samples. Included in these analyses were: New Guinea Highlanders as a source of Papuan ancestry; Han Chinese as a source of Asian ancestry; Kankanaey as a source of Austronesian ancestry; Tolai (mixed Papuan/Austronesian ancestry) and Baining_Marabu (Papuan ancestry only) from New Britain to investigate relationships with the Bismarck Archipelago; modern Vanuatu with mixed Papuan/Austronesian ancestry; and the ancient Guam, Lapita Vanuatu and Lapita Tonga samples. We also included Mbuti as an outgroup. We first constructed a maximum-likelihood tree and added migration edges, using the software TreeMix (58); a tree with 2 migration edges (Figure 5A) has all residuals within 3 SE (Fig. S12) and thus provides a reasonable fit. This tree indicates shared drift between the ancient Guam and Lapita samples, with the migration edges bringing Lapita-related ancestry into the modern Vanuatu and Tolai samples.

**Figure 5.**
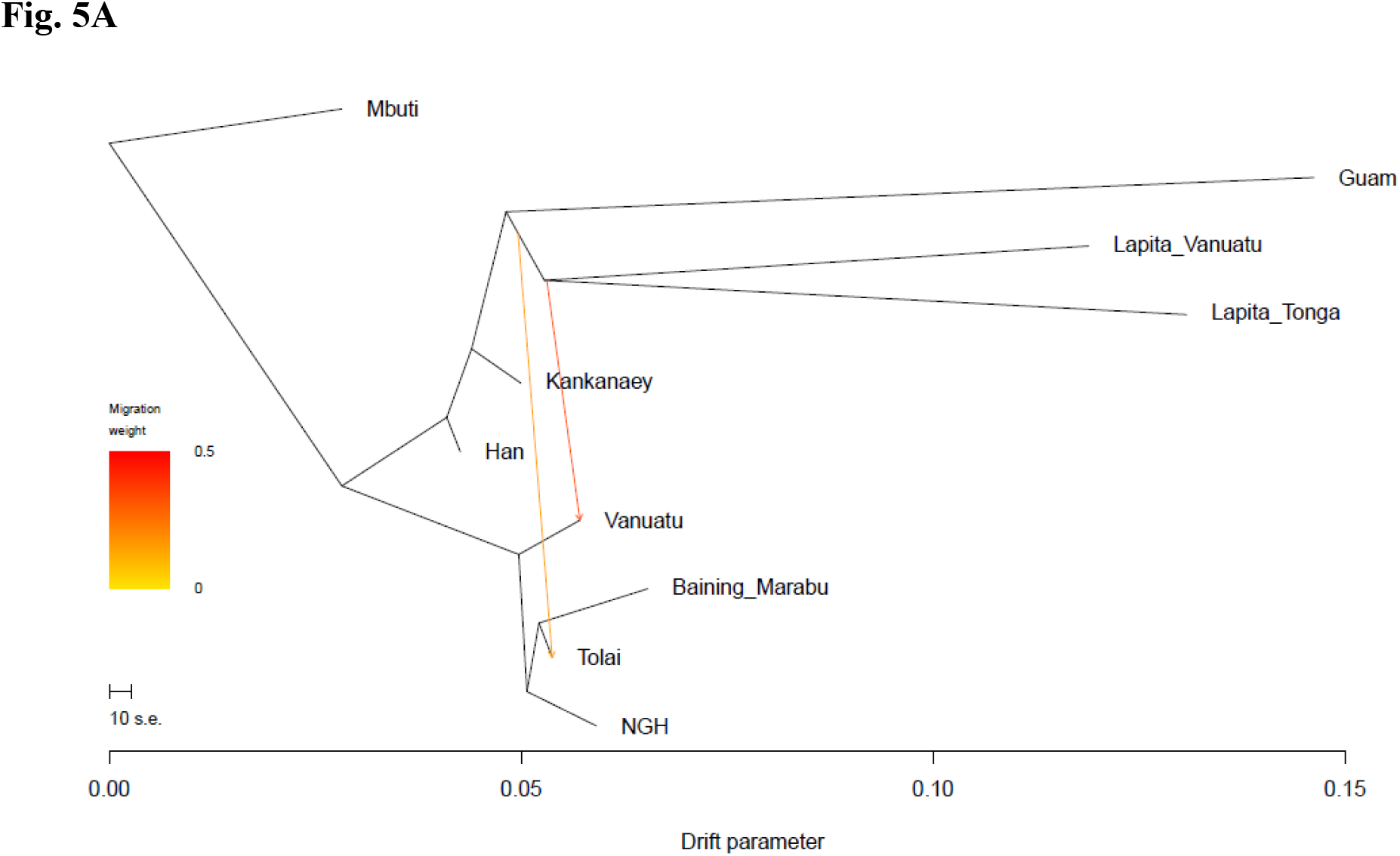
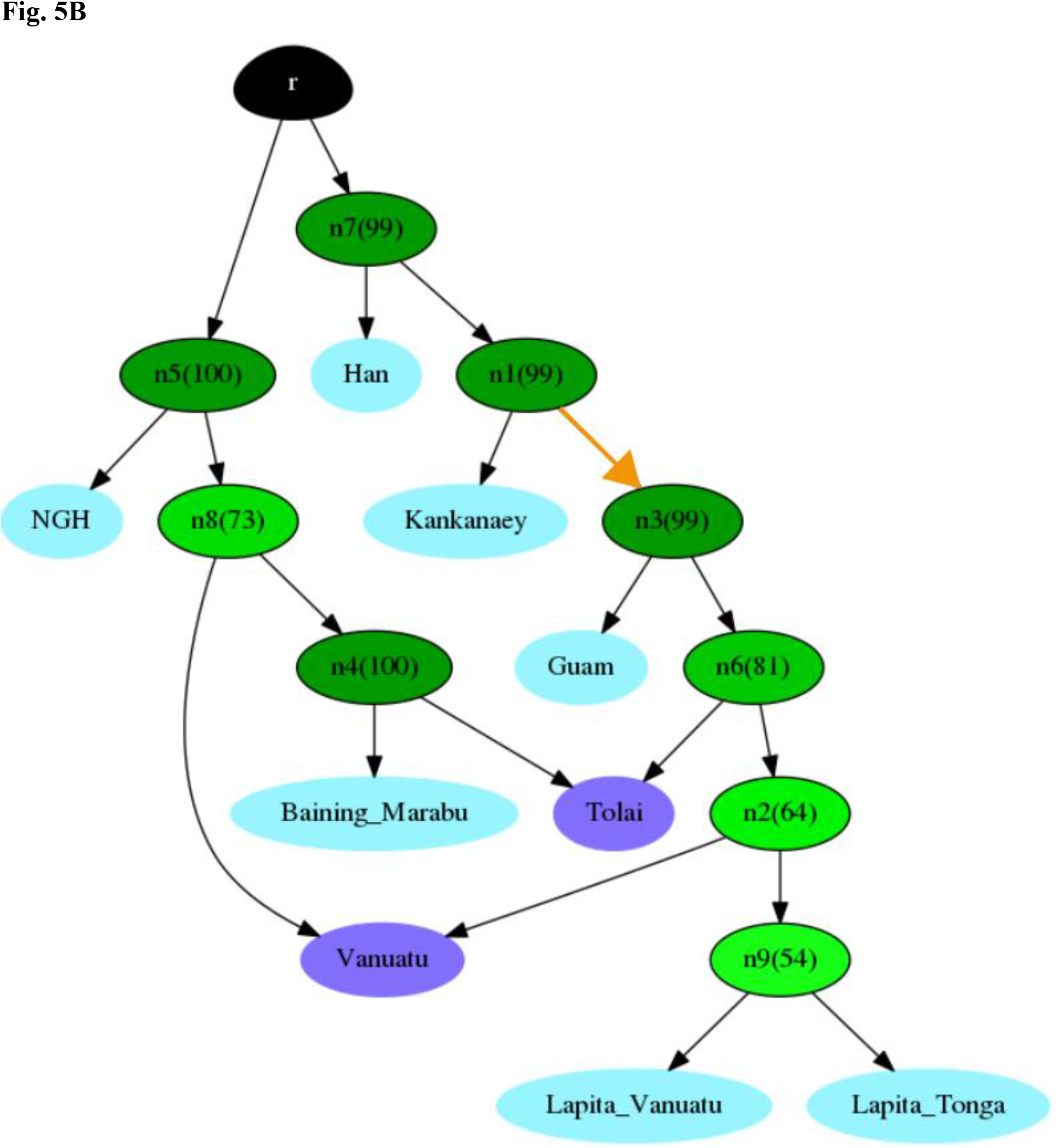
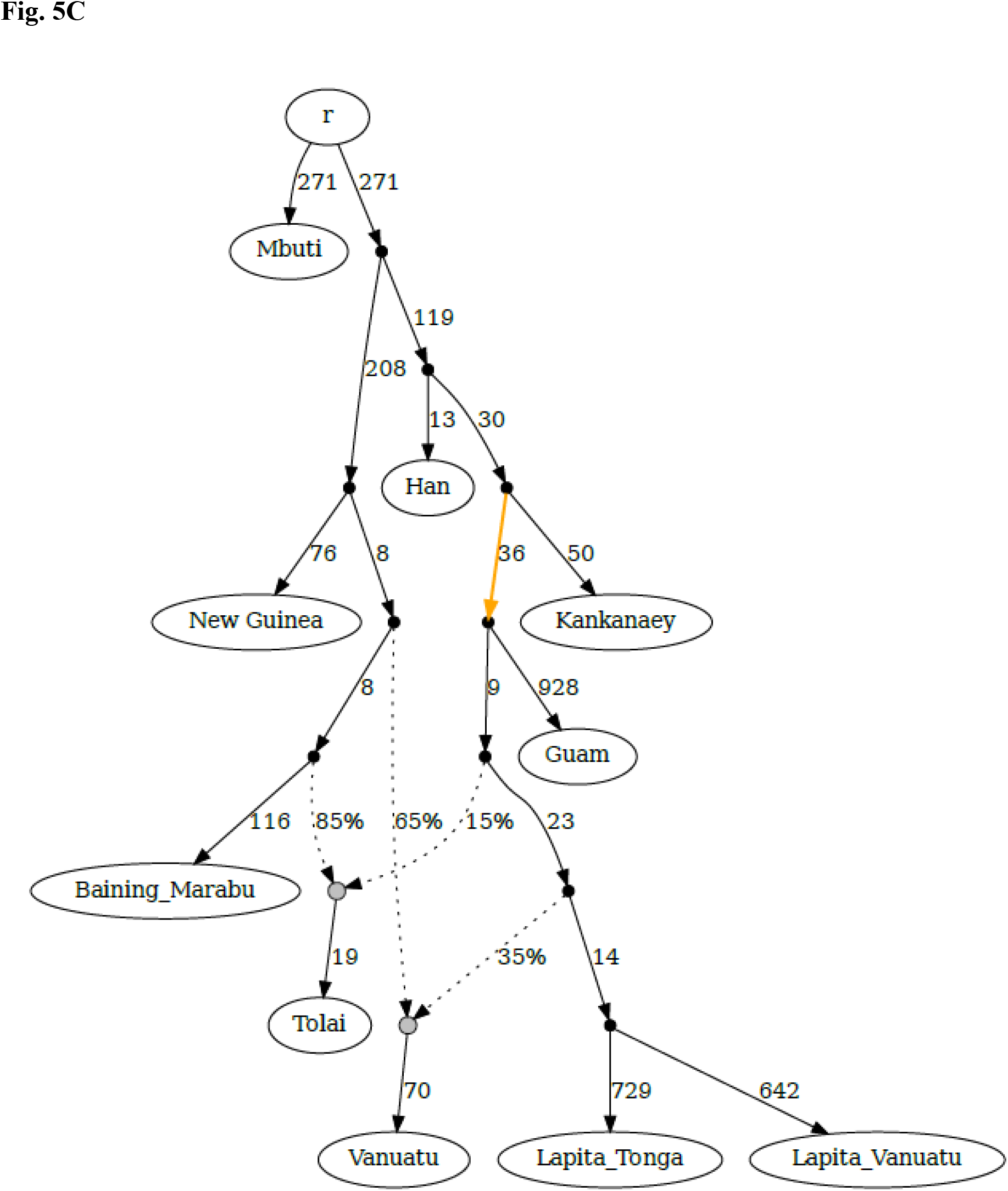
Tree and graph depictions of the relationships of ancient Guam, early Lapita, and select Asian and Oceanian populations. **A**, maximum-likelihood tree with two migration edges. All residuals (Fig. S12) are within 3 SE. **B**, consensus graph with nodes present in at least 50% of the topology sets recovered with AdmixtureBayes. **C**, admixture graph obtained with qpGraph for the topology found by AdmixtureBayes with the highest posterior probability (Fig. S13). The colored arrows in **B** and **C**indicate drift (ancestry) shared between the ancient Guam and early Lapita samples.

We additionally investigated admixture graphs using a Markov Chain Monte Carlo method, implemented in the software AdmixtureBayes (59), to sample the space of possible admixture graphs. The graph with the highest posterior probability (17.6%) supports shared drift between the ancient Guam and early Lapita samples (Fig. S13); moreover, a consensus graph that depicts the nodes present in at least 50% of the 1000 graphs with the highest posterior probabilities (Figure 5B) indicates that the shared drift between the ancient Guam and early Lapita samples (node n3 in Figure 5B) appears in 99% of the topologies. We further examined this topology, inferred in an unsupervised manner by both TreeMix and AdmixtureBayes, with a combination of *f*-statistics using the qpGraph software. This topology has a worst-fitting Z score of 4.56 (Figure 5C), which is above the conventional threshold of the worst-fitting |Z-score| <3 for an “acceptable” graph. Deviations between the fitted and observed data can be explained either by an incorrect topology (which, in the case of qpGraph, is specified by the user and not inferred from the data) or by unmodeled admixture. The worst-fitting *f-*statistics tend to involve Han Chinese; when they are excluded the worst-fitting Z score is reduced to −3.72. This graph has five *f-*statistics with |Z-score| >3, all of which involve Mbuti and New Guinea Highlanders, so this graph probably provides a reasonable depiction of the relationships of the Oceanian samples, in particular the shared drift between the ancient Guam and early Lapita samples. For the two populations with mixed ancestry, the modern Vanuatu sample is inferred to have 65% Papuan-related and 35% Austronesian-related ancestry, while the Tolai sample has 85% Papuan-related and 15% Austronesian/Lapita-related ancestry; these estimates are in close agreement with those from AdmixtureBayes (Vanuatu: 66% Papuan-related and 34% Austronesian-related ancestry; Tolai: 87% Papuan-related and 13% Austronesian-related ancestry).

We further investigated the shared drift between the ancient Guam and early Lapita samples by including ancient samples from Liangdao that share ancestry with aboriginal Taiwanese (60) in the admixture graph analyses. While the results suggest that Liangdao is a better proxy than modern samples for the Austronesian-related ancestry in the ancient Guam and early Lapita samples (Fig. S14), there is still shared drift between the ancient Guam and early Lapita samples.

## Discussion

Some caution is warranted in interpreting the results of this study of ancient DNA from Guam, as they are based on two skeletons that may be related and that date from approximately 1400 years after the first human settlement of Guam. Previous studies of ancient DNA from Remote Oceania have found that initial results based on a limited number of samples (25) did not capture the full complexity revealed when additional samples were analyzed (23, 24). Nonetheless, the relationships that these ancient Guam samples exhibit with other ancient samples, as well as with modern samples from the region, provide some interesting insights into the peopling of Guam and the further settlement of Remote Oceania that should be the basis for further investigations.

### Origins of the ancient Guam samples

The mtDNA and Y chromosome haplogroups of the ancient Guam samples suggest links with Southeast Asia rather than New Guinea or the Bismarck Archipelago. Moreover, none of the analyses of the genome-wide data found any trace of Papuan-related ancestry in the ancient Guam samples. Our results thus rule out any source for the ancestry of these individuals that is east of Wallace’s line, as substantial amounts of Papuan-related ancestry are present in eastern Indonesia, New Guinea, and the Bismarck Archipelago. The most likely source is the Philippines, although western Indonesia is also possible; further sampling of Philippine and Indonesian populations – and ancient DNA from these regions – would help pinpoint the source. Moreover, in considering the archaeological evidence, finer-scale sampling is needed to contend with a rapid geographic spread of the red-slipped pottery horizon around 3.5 kya, reflecting population dispersal from the Philippines both eastward into the Marianas and southward into Sulawesi, as well as eventually further.

A Philippine source for the foundational population of Guam is consistent with the findings of modern DNA sampling (27), the linguistic evidence (1), and the archaeological signature at the time of first Marianas settlement about 3.5 kya (29, 42). However, computer simulations of sea voyaging instead have indicated New Guinea or the Bismarck Archipelago as probable origin points of voyages reaching the Marianas (32, 33). One potential scenario to reconcile these two lines of evidence is that people traveled from the Philippines to New Guinea or the Bismarcks, without mixing with any populations along the way, and then voyaged from New Guinea/the Bismarcks to Guam, again without first mixing with any resident populations. However, the TreeMix and AdmixtureBayes results (Figure 5) do not support this scenario, nor does the linguistic and archaeological evidence. In particular, the earliest pottery in the Marianas, dating to around 3.5 kya (2, 42), likely predates the oldest Lapita sites to the east of New Guinea, dated to not more than 3.3 kya (4). Yet the pottery, fine shell ornaments, and other cultural objects in the Marianas dating to 3.5 kya are quite distinct from the Lapita tradition, and instead can be linked to material markers in the Philippines that date to 3.8-3.5 kya (18, 29, 61), thus supporting movement from the Philippines to the Marianas. Moreover, the computer simulations of sea voyaging do not adequately consider the ability of ancient voyagers to travel against strong ocean currents and prevailing winds; in particular, the single outrigger canoes of the Chamorros - the ‘flying proas’ - impressed early visitors with their greater speed and maneuverability, compared to Spanish ships (62). There is even at least one historically documented event of a Chinese trader drifting in a “sampan” from Manila to Guam during the 1600s (63). Ancient DNA from early Lapita skeletons in the Bismarcks would provide a further test of the hypothesis that people moved from the Bismarcks to Guam.

### Relationships between ancient Guam and early Lapita samples

> “What about a Micronesian route [for the colonization of Polynesia]? It is not in favor with the anthropologists, though after all it was not anthropologists who settled Polynesia.”
>
> — -William Howells, *The Pacific Islanders* (1973), p. 253

All analyses consistently point to a surprisingly close relationship between the ancient Guam and early Lapita samples. This closeness is particularly evident in the outgroup *f*3 and various *f*4 analyses (Figure 4, Figs. S9-S11), and in the TreeMix and admixture graph results (Figure 5), all of which indicate shared ancestry between the ancient Guam and early Lapita samples. Moreover, admixture graphs indicate that the ancient Guam samples diverged first, and do not support movement of people from the Bismarcks to Guam (Figure 5; Figs. S13- S14). However, the admixture graph results should be viewed with caution, as they may be influenced by including a mix of ancient and modern DNA samples in the analyses (usually with fewer ancient than modern samples for each population), with possible attractions between ancient samples due to similar patterns of contamination and/or sequencing errors due to damage. Nonetheless, it appears that people either moved from the Marianas to the Bismarcks and then to Remote Oceania, or that the ancestors of the ancient Guam and early Lapita samples migrated separately, and by different routes, from the same source population.

Our results do not allow us to distinguish between these two possibilities. Nevertheless, we point out that a direct movement of people from the Philippines (or nearby areas) to the Bismarcks, either via the Marianas or by some other path that bypassed eastern Indonesia and the rest of New Guinea, would account for one peculiar observation, and that is the lack of Papuan-related ancestry in the early Lapita samples from Vanuatu and Tonga (23–25). If the ancestors of Polynesians migrated from Taiwan or the Philippines to the Bismarcks by island-hopping through eastern Indonesia and along the coast of New Guinea (Figure 1), in a process that took a few hundred years (perhaps 10-15 generations), then they would have encountered people with Papuan-related ancestry along the way, and there would have been ample opportunity for them to have picked up some Papuan-related ancestry. Perhaps the ancestors of Polynesia did move via this route, but did not immediately mix with the people along the way, because of social or other perceived differences. However, any such barrier to mixing did not last long, as Papuan-related ancestry shows up in Vanuatu almost at the same time as the early Lapita samples (24), and there is evidence for substantial later Papuan-related contact in Vanuatu, Santa Cruz, and Fiji (23, 24, 26, 36). An alternative explanation that is worth considering is that the early ancestors of Polynesians lack Papuan-related ancestry because they did not encounter people with Papuan-related ancestry until they reached the Bismarcks – perhaps because they voyaged via the Marianas or otherwise bypassed eastern Indonesia and coastal New Guinea.

As the quotation from Howells (1973) at the beginning of this section indicates, the settlement of Polynesia via Micronesia has generally not been considered by researchers. However, this possibility has been suggested based on pottery evidence (17), and the genetic evidence presented here provides further support, as well as additional insights into the connections between Micronesians and Polynesians noted previously (11–16). Howell’s suggestion of a role for Micronesia (specifically, the Marianas) in the settlement of Polynesia merits further consideration.

## Methods

### Site description and samples

The two skeletons RBC1 and RBC2 were uncovered outside Ritidian Beach Cave (also called Ritidian First Cave), within the larger Ritidian Site of northern Guam (Figs. S1 and S2). The two individuals had been buried side by side, in extended position inside distinctive pits. The heads and torsos had been removed slightly later. Details of these findings have been reported elsewhere and situated within the larger site chronology and context (43). The two skeletons from Ritidian offer a rare view of ancient burial practice in the Marianas region, as similar burial practices have been observed in the Philippines (64, 65) and Indonesia (66). While the site and indeed this specific cave revealed multiple cultural occupation layers dating back to the first regional settlement about 3.5 kya, these two burials of RBC1 and RBC2 were found within the layer of approximately 2.5 – 2 kya, confirmed by direct radiocarbon date from a bone of RBC2 of 2180 +/- 30 ybp (43). A tarsal bone was provided from each skeleton for ancient DNA analysis.

### DNA extraction, library preparation, and whole-genome sequencing

In an ancient DNA clean room, approximately 1 mm of material was removed from the surface of each specimen and ~50 mg bone powder obtained by drilling into the bone with a dentistry drill at low speed. DNA was extracted following a protocol provided elsewhere, using spin columns and binding buffer option ‘D’ (67). DNA libraries were prepared from 10-μl aliquots of each DNA extract using an automated protocol for single-stranded library preparation (68) with a Bravo NGS workstation. Negative controls were included both during DNA extraction and library preparation; these contained water instead of sample powder or DNA extract, respectively. The number of library molecules obtained from each sample DNA extract was more than 100 times higher than in the extraction and library negative controls (Table S1). All libraries, including the negative controls, were then amplified and double-indexed (69) as described elsewhere (68).

Whole-genome sequencing data were generated on the Illumina HiSeq 2500 platform (2x 76bp paired-end sequencing). After de-multiplexing (requiring a perfect index), overlapping paired-end sequences were merged into full-length molecule sequences (70) and subsequently aligned to the human reference genome *hg19* with decoy sequences (ftp://ftp-trace.ncbi.nih.gov/1000genomes/ftp/technical/reference/phase2_reference_assembly_sequence/hs37d5.fa.gz), using bwa aln(71) with parameters optimized for ancient DNA (“-n 0.01 -o 2 -l 16500”(72)). The sequencing data were filtered for a minimum read length of 35 bp and a minimal mapping quality of 25. Duplicate reads were removed using DeDup(73) and the number of substitutions compared to the human reference genome was quantified using damageprofiler (https://github.com/Integrative-Transcriptomics/DamageProfiler). Finally, we subset the sequencing data to reads for which we observed a C-to-T substitution in the first three bases at either read end (Table S1).

### MtDNA enrichment and sequencing

Libraries were enriched for human mitochondrial DNA using a synthetic probe set (74) encompassing the revised Cambridge reference sequence (rCRS) (75) in 1-bp tiling. Hybridization capture was performed in two successive rounds, following an on-bead capture protocol (76) implemented on the Bravo NGS workstation (77). The enriched libraries were pooled with libraries from other projects and sequenced on an Illumina MiSeq in paired-end mode (2x 76 cycles).

The sequencing data were processed as described above for the whole-genome sequencing data, but mapped to the rCRS using bwa aln with the same settings. Sequences were assigned to their respective source libraries, requiring perfect matching of both indices. Sequences that were shorted than 35 bp or that did not produce alignments with a map quality of at least 25 were discarded. PCR duplicates were removed using DeDup (73). After discarding libraries with contamination > 25% (Table S2), estimated by a likelihood-based method (46), we obtained 32,386 unique reads for RBC1 and 94,116 unique reads for RBC2 (Table S3). Elevated frequencies of C to T substitutions at the beginning and end of sequence alignments, which result from cytosine deamination in ancient DNA (78, 79), were detected in the mtDNA reads (Table S2).

An in-house pipeline (https://github.com/alexhbnr/mitoBench-ancientMT) was used to call the mtDNA consensus sequence, which required a minimum of 3 reads and used snpAD (80) to infer the consensus allele while taking into account ancient DNA damage. Contamination was estimated by a likelihood-based method (46), and HaploGrep2 (81) was used to call mtDNA haplogroups.

### Genome-wide SNP capture enrichment and sequencing

Enrichment of the libraries for a panel of approximately 1.2 million SNPs was performed using a set of DNA capture probes (“1240k”, composed of SNP panels 1 and 2 as described elsewhere (44)) and two successive rounds of in-solution hybridization capture (74). Sequencing of the enriched libraries and raw data processing were performed as described for the whole-genome sequencing above. Genotypes were inferred by randomly sampling an allele observed at each site after masking Ts at the five terminal bases at each read end by replacing them with Ns. For determining the Y chromosome haplogroup of the male sample, we subset the genotypes to the sites located on the human Y chromosome and analyzed them using yHaplo (82) with the non-default option *“--ancStopThresh 1e6”.*

The sequencing data have been made available at the European Nucleotide Archive (ENA) under the accession id PRJEB40707.

### Genome-wide SNP data analysis

#### Comparative datasets

Newly generated data from Guam were merged with published data from modern and ancient samples (Tables S7 and S8) as follows. First, for comparisons to populations from Island Southeast Asia, the Guam data were merged with previously-curated data from 25 modern populations genotyped on the Affymetrix 6.0 array (26, 83, 84); to provide worldwide context these data were further merged with a subset of the whole genome sequences from the Simons Genome Diversity Project (SGDP; (85)). Related individuals were identified based on kinship coefficients, estimated using the software KING (86), with subsequent removal of one individual from each pair. Pruning of SNPs in LD was done using the PLINK tool (86) with the following settings: -indep- pairwise 200 25 0.4 (87). After these quality filtering steps, there were 136,162 SNPs and 305 individuals from 51 populations from Eurasia and Oceania remaining for the analyses. This dataset was used only for PCA and ADMIXTURE analyses.

Second, to better resolve relationships with populations from Near and Remote Oceania, as well as with other ancient samples from Asia and the Pacific, we used data from 53 modern populations from Oceania and 39 populations from East Asia genotyped on the Affymetrix Human Origins array (23, 25, 88–90), as well as previously published shotgun and capture-enrichment sequencing data from 82 ancient samples (23–25, 45, 60, 91). After removing related individuals as described above for the Affymetrix 6.0 data, this dataset consisted of 1194 individuals and 593,124 SNPs. Not all samples were used for all of the analyses. For PCA and ADMIXTURE analyses, we used an LD-pruned dataset of 216,996 SNPs. In addition, ancient samples with more than 15,000 missing sites were excluded from the ADMIXTURE analysis.

#### Data Analyses

We attempted to estimate relatedness between RBC1 and RBC2 by calculating the fraction of pairwise differences at 33,040 overlapping sites that are included on the Human Origins array. For comparison, we also calculated this fraction for modern samples from the 1000 Genomes Project data set (56), which includes individuals with known degrees of relatedness, and for ancient samples from Southeast Asia and Oceania; the ancient DNA data were obtained from the Reich lab website (https://reich.hms.harvard.edu/downloadable-genotypes-present-day-and-ancient-dna-data-compiled-published-papers; v42.4).

Principle components analysis was performed as described previously (92) with one modification, namely for the analyses which included ancient samples, the principle axes were calculated based on modern samples, and the ancient samples were projected using least-squares projection (which is more appropriate than orthogonal projection for samples with high amounts of missing data), as described in the documentation to the smartpca software (93).

To infer individual ancestry components and analyze population structure, we used the ADMIXTURE software (94) in the unsupervised mode. For each dataset, we first removed SNPs in strong LD (r2 > 0.4) using the PLINK tool (95), and for the HO dataset we further excluded ancient samples which had less than 15,000 SNPs remaining. We varied the number of ancestral populations (K value) from K=2 to K=8 for the Affymetrix 6.0 dataset, and from K=5 to K=12 for the Human Origins Array dataset. We performed 100 independent runs for each value of K, and used the cross-validation procedure implemented in the ADMIXTURE software to assess the best value of K.

To formally test population relationships suggested by PCA and ADMIXTURE analyses, we used outgroup-*f*3 and *f*4-statistics, implemented in the ADMIXTOOLS software suite (89). All data processing and analyses were carried out using the admixr R package (96).

To model the relationships between modern and ancient samples, we first used the unsupervised TreeMix (58) and AdmixtureBayes methods (59) to infer topologies, that were then tested using the qpGraph software implemented in ADMIXTOOLS (89). We performed ten independent runs of TreeMix with 0 to 5 migration events, and report the tree with the highest likelihood. For the AdmixtureBayes analyses we increased the default number of MCMC steps to 1,000,000, as recommended by the developers to avoid convergence problems for a model with 10 populations. We used the ten topologies with the highest posterior probabilities estimated by AdmixtureBayes as input graphs for qpGraph, which we ran with parameters: blgsize: 0.05, forcezmode: YES, lsqmode: YES, diag: .0001, bigiter: 6, hires: YES, lambdascale: 1. All three methods were applied to the exact same dataset, using all samples available for each population (Supplementary Table 7), with the exception of the admixed modern Vanuatu. The amount of Polynesian ancestry in this population is highly variable, with a range of 9%-38% today, so for our analyses we took all individuals from the island of Futuna, where Polynesian-related ancestry is highest (23). TreeMix and AdmixtureBayes do not allow sites with missing data, so for each SNP each population is required to have at least one genotype call. Since our model included three ancient populations, the number of sites available for these analyses was reduced to 76,284. For qpGraph it is possible to use an option which would maximise the number of sites for each computed statistic, but since we have modern and ancient data in the same analysis, this would result in dramatically uneven SNP sets for different comparisons. As this could bias the results, we therefore chose not to use this option. Mbuti was used as an outgroup in all of the admixture graph analyses.

Statistical programming was done using the statistical program R version 4.0.1 (https://www.R-project.org/). We used the tidyverse (97), data.table (https://CRAN.R-project.org/package=data.table), Hmisc (https://CRAN.R-project.org/package=Hmisc), and pheatmap (https://CRAN.R-project.org/package=pheatmap) packages.

## Acknowledgements

We thank S. Nagel, B. Nickel, B. Schellbach, A. Schmidt, A. Weihmann and M. Wunsch for performing the laboratory work, the Bioinformatics Group in the Dept. of Evolutionary Genetics at the Max Planck Institute for Evolutionary Anthropology for the initial processing of the sequencing data, P. Bellwood and R. Blust for comments on an earlier draft, and M. Hajdinjak for helpful discussion. This particular archaeology study at the Ritidian Site in Guam was permitted and coordinated with the U.S. Fish and Wildlife Service, and it was funded by the Chiang Ching-kuo Foundation (grant number: RG021-P-10) and by the Australian Research Council (grant number: DP150104458). Research was funded by the Max Planck Society.

## Notes

### Competing Interest Statement

The authors have declared no competing interest.

